# How accurate are citations of frequently cited papers in biomedical literature?

**DOI:** 10.1101/2020.12.10.419424

**Authors:** V Pavlovic, T Weissgerber, D Stanisavljevic, T Pekmezovic, V Garovic, N Milic, CITE Investigators

**Author notes:** **Corresponding author:** Natasa Milic, MD, PhD.

## Abstract

Citations are an important, but often overlooked, part of every scientific paper. They allow the reader to trace the flow of evidence, serving as a gateway to relevant literature. Most scientists are aware of citations errors, but few appreciate the prevalence or consequences of these problems. The purpose of this study was to examine how often frequently cited papers in biomedical scientific literature are cited inaccurately. The study included an active participation of first authors of frequently cited papers; to first-hand verify the citations accuracy. The approach was to determine most cited original articles and their parent authors, that could be able to access, and identify, collect and review all citations of their original work. Findings from feasibility study, where we collected and reviewed 1,540 articles containing 2,526 citations of 14 most cited articles in which the 1^st^ authors were affiliated with the Faculty of Medicine University of Belgrade, were further evaluated for external confirmation in an independent verification set of articles. Verification set included 4,912 citations identified in 2,995 articles that cited 13 most cited articles published by authors affiliated with the Mayo Clinic Division of Nephrology and Hypertension (Rochester, Minnesota, USA), whose research focus is hypertension and peripheral vascular disease. Most cited articles and their citations were determined according to SCOPUS database search. A citation was defined as being accurate if the cited article supported or was in accordance with the statement by citing authors. A multilevel regression model for binary data was used to determine predictors of inaccurate citations. At least one inaccurate citation was found in 11% and 15% of articles in the feasibility study and verification set, respectively, suggesting that inaccurate citations are common in biomedical literature. The main findings were similar in both sets. The most common problem was the citation of nonexistent findings (38.4%), followed by an incorrect interpretation of findings (15.4%). One fifth of inaccurate citations were due to “chains of inaccurate citations,” in which inaccurate citations appeared to have been copied from previous papers. Reviews, longer time elapsed from publication to citation, and multiple citations were associated with higher chance of citation being inaccurate. Based on these findings, several actions that authors, mentors and journals can take to reduce citation inaccuracies and maintain the integrity of the scientific literature have been proposed.

## Introduction

Citations are an important, but often overloooked, part of every scientific paper. They allow the reader to trace the flow of ideas and evidence through a paper, serving as a gateway to other relevant literature. Citations also allow readers to confirm that the cited information supports the authors’ hypotheses and suppositions. Many deficits in citation practices unfortunately have been reported. A recent meta-analysis demonstrated that 25.4% of papers contained a citation error (1). Van der Vet and Nijveen (2) reported that even retracted articles continue to be cited approvingly years after they have been retracted. Citation errors can have serious implications. Millions of Americans suffer from opioid addiction and opioids contributed to 47,600 drug overdose deaths in the United States in 2017 (3). A team of Canadian researchers proposed that uncritical citations of a letter published in the New England Journal of Medicine (4) may have contributed to the opioid crisis (5). The letter stated that narcotic addiction was rare in hospital inpatients with no histories of addiction (4). This five sentence letter contained no detailed methods or results (4); yet it was cited hundreds of times as evidence that addiction risk was low when opioids were perscribed for chronic pain (5). Some citations clearly distorted the findings of the letter, and 81% of the letter’s 608 citations did not mention that the study only included hospital inpatients. Leung et al. (5) argue that these uncritical and misleading citations may have helped to shift perscribing practices by convincing doctors that addiction risk was low with chronic opioid use.

A published letter attempting to correct an overestimate of the number of Cochrane reviews on rehabilitation interventions (6) provides another example of the dangers of citation copying (7). Although the article (6) was cited 62 times, all of these citations were related to meta-analyses of genetic risk factors (7). Authors cited the letter to support their use of the Cochran Q-statistic for exploring heterogeneity of effect sizes, whereas the letter was written to highlight the need for Cochrane reviews on rehabilitation.

While the academic literature is replete with similar errors, assessing citation accuracy takes considerable effort. Most scientists are aware of citation errors and copying, but few appreciate their prevalence or consequences. The purpose of this study was to: 1) examine how often frequently cited papers in biomedical literature are cited inaccurately, 2) explore factors associated with inaccurate citations and 3) discuss actions that authors, mentors, and journals can take to eliminate citation errors.

## Methods

The study was designed to include an active participation of first authors of the frequently cited papers in biomedical scientific literature, to first-hand verify the accuracy of the citations of their original work. The approach was to determine most cited original articles and their parent authors, which could be feasible to access, and identify, collect and review all citations of their original work throughout the biomedical scientific literature. As this approach resulted in a time consuming project that is complex to manage, we had conducted a feasibility study, whose results were than further evaluated for external confirmation in an independent verification set of articles. The study was approved by the Ethical Committees of the Faculty of Medicine University of Belgrade (2650/IV-6) and the Mayo Clinic, Rochester, MN, USA (19-005085).

### Feasibility study

The sample was formed in two stages (Figure 1). We first chose “source articles” - the most cited articles in which the 1^st^ authors were affiliated with the Faculty of Medicine University of Belgrade, according to a SCOPUS bibliographic database search on October 1, 2017. Based on our hypothesis that the frequency of citation inaccuracies would be 10%, we calculated that 1500 citing articles would be needed to estimate the frequency of inaccurate citations with a precision of 1.5% (alpha=0.05). Fourteen source articles were chosen to reach this predetermined sample size (n=1500 citing articles). Characteristics of the source articles, including the number of citations, are presented in Table 1. Source articles were published between 1994 and 2009. The time elapsed from the publication of the article to the beginning of this study was between 8 and 23 years, and the total number of source article citations for this period ranged from 63 to 393. The publication field was determined according to journal classification from Journal Citation Reports (JCR). In the second stage, we collected all “citing articles” which cited the source articles (according to a SCOPUS bibliographic database search on October 1, 2017). Citing articles were included in the study if they were published in English and we could obtain a full text version of the article. Articles written in other languages, as well as books and book chapters were excluded. The final sample included 1540 of the 1565 citing articles published in English; 25 citing papers (1.6%) were excluded because a full-text version of the manuscript could not be retrieved.

**Table 1.** Characteristics of source articles in the feasibility study

| Source article | Publication field | Publication year | Years since publication | Number of citations |  |  |  |
| --- | --- | --- | --- | --- | --- | --- | --- |
|  |  |  |  | Articles in English | Articles not in English* | Books and book chapters* | Total citations |
| 1 | Multidisciplinary Sciences | 1997 | 20 | 353 | 16 | 24 | 393 |
| 2 | Cardiac & Cardiovascular Systems, Peripheral Vascular Disease | 1994 | 23 | 184 | 35 | 15 | 234 |
| 3 | Endocrinology & Metabolism | 1995 | 22 | 179 | 5 | 1 | 185 |
| 4 | Rheumatology | 2009 | 8 | 127 | 3 | 3 | 133 |
| 5 | Infectious Diseases | 2005 | 12 | 102 | 8 | 7 | 117 |
| 6 | Endocrinology and Metabolism, Nutrition & Dietetics | 2009 | 8 | 108 | 1 | 4 | 113 |
| 7 | Clinical Neurology, Psychiatry, Surgery | 1994 | 23 | 85 | 18 | 9 | 112 |
| 8 | Clinical Neurology | 2002 | 15 | 76 | 9 | 4 | 89 |
| 9 | Clinical Neurology | 2001 | 16 | 56 | 21 | 4 | 81 |
| 10 | Urology & Nephrology | 1996 | 21 | 45 | 31 | 2 | 78 |
| 11 | Clinical Neurology | 2001 | 16 | 71 | 4 | 1 | 76 |
| 12 | Medicine, Legal, Pathology | 2003 | 14 | 61 | 6 | 1 | 68 |
| 13 | Immunology, Neurosciences | 2001 | 16 | 62 | 2 | 3 | 67 |
| 14 | Peripheral Vascular Disease | 2006 | 11 | 56 | 4 | 3 | 63 |
| Total |  |  |  | 1,565 | 163 | 81 | 1,809 |

\* These categories were excluded from subsequent analyses

**Figure 1.**
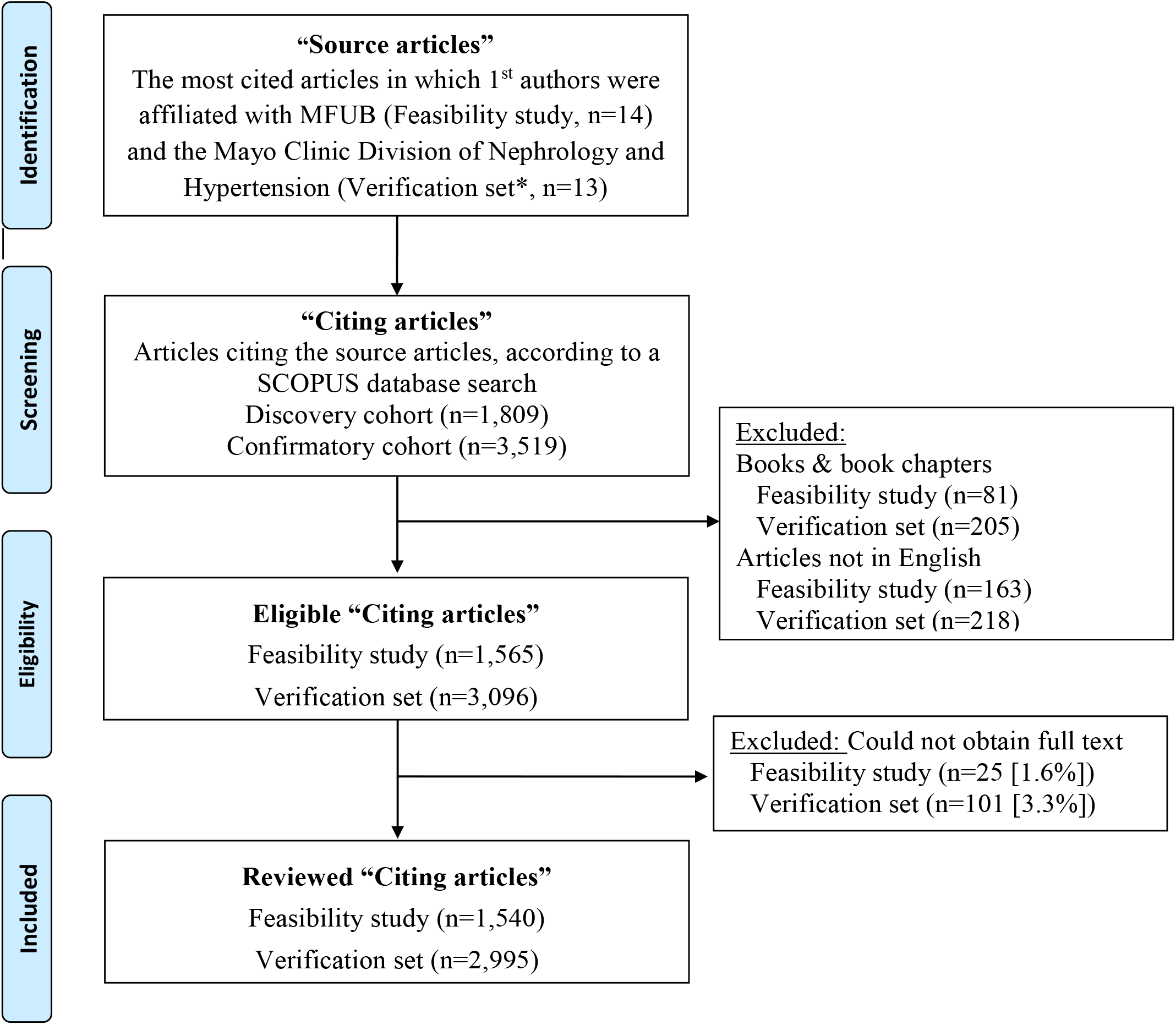
Study design flow chart Abbreviation: MFUB, Medical Faculty University of Belgrade *two authors did not agree to participate and were replaced with next authors from the list

### Assessing Citation Accuracy

A citation was defined as being accurate if the reference (source article) supported or was in accordance with the statement by citing authors. Each citing article was first reviewed for citation accuracy and discussed by three reviewers. If inaccuracies were detected, citations were further evaluated by one of the authors of the source article, including first authors (n=11) or another co-author and active member of the research team (n=3). Each author checked the accuracy of citations of his or her own paper and classified the type of inaccurate citation as follows: citation of nonexistent findings, incorrect interpretation of findings, incorrectly cited method, incorrectly cited numerical data/results, citation of nonexistent numerical data/results, wrong context, cited findings from another source, or reference listed in the bibliography but not cited in the text. We extracted the following data from each citing article to identify factors associated with citation inaccuracies: publication year, article type as defined by the journal in which the article was published (i.e. original article, review, perspective, editorial, etc.), self-citation, referencing style, number of authors, impact factor of the journal, and number of references in the bibliography. The impact factor was extracted from Journal Citation Reports (JCR), if available, for the year in which the citing paper was published. Citation was considered as a self-citation if a citing paper and source paper had at least one author in common.

### Verification set

Source articles were the most cited articles published by authors affiliated with the Mayo Clinic Division of Nephrology and Hypertension (Rochester, Minnesota, USA), whose research focus is hypertension and peripheral vascular disease. Most cited articles were determined by a SCOPUS database search on May 1, 2019. It was planned to include twice as many citing articles in the verification set compared to the feasibility study. The final verification sample included 2,995 of the 3,096 citing articles published in English; 3.3% of articles were excluded due to unavailability of a full-text version of the manuscript. The procedure of reviewing citing articles was the same as in the feasibility study. Characteristics of the source articles in the verification set are presented in Table 2.

**Table 2.**
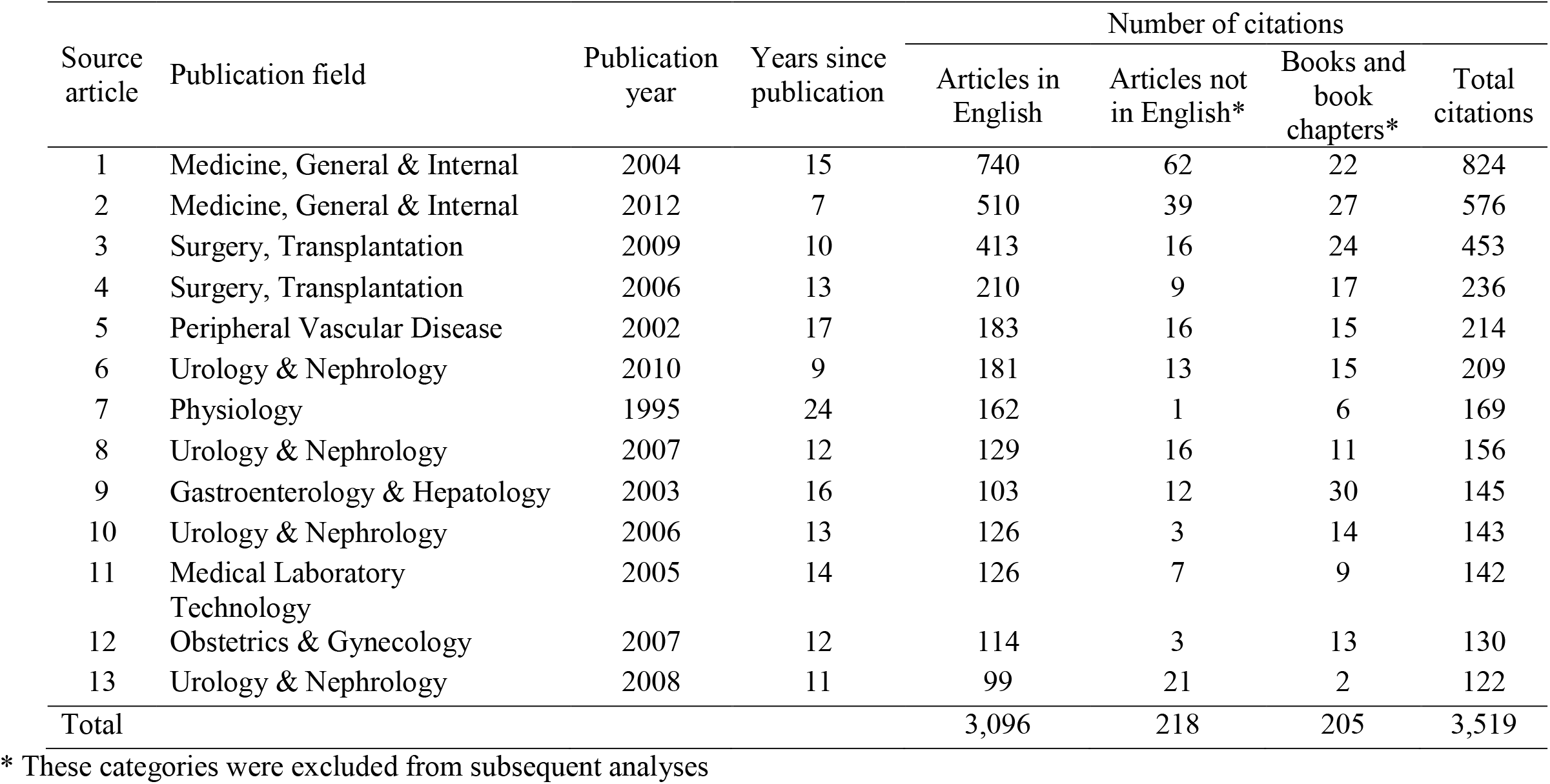
Characteristics of source articles in the verification set

| Source article | Publication field | Publication year | Years since publication | Number of citations |  |  |  |
| --- | --- | --- | --- | --- | --- | --- | --- |
|  |  |  |  | Articles in English | Articles not in English* | Books and book chapters* | Total citations |
| 1 | Medicine, General & Internal | 2004 | 15 | 740 | 62 | 22 | 824 |
| 2 | Medicine, General & Internal | 2012 | 7 | 510 | 39 | 27 | 576 |
| 3 | Surgery, Transplantation | 2009 | 10 | 413 | 16 | 24 | 453 |
| 4 | Surgery, Transplantation | 2006 | 13 | 210 | 9 | 17 | 236 |
| 5 | Peripheral Vascular Disease | 2002 | 17 | 183 | 16 | 15 | 214 |
| 6 | Urology & Nephrology | 2010 | 9 | 181 | 13 | 15 | 209 |
| 7 | Physiology | 1995 | 24 | 162 | 1 | 6 | 169 |
| 8 | Urology & Nephrology | 2007 | 12 | 129 | 16 | 11 | 156 |
| 9 | Gastroenterology & Hepatology | 2003 | 16 | 103 | 12 | 30 | 145 |
| 10 | Urology & Nephrology | 2006 | 13 | 126 | 3 | 14 | 143 |
| 11 | Medical Laboratory Technology | 2005 | 14 | 126 | 7 | 9 | 142 |
| 12 | Obstetrics & Gynecology | 2007 | 12 | 114 | 3 | 13 | 130 |
| 13 | Urology & Nephrology | 2008 | 11 | 99 | 21 | 2 | 122 |
| Total |  |  |  | 3,096 | 218 | 205 | 3,519 |

\* These categories were excluded from subsequent analyses

### Statistical analysis

Descriptive statistics, including numbers and percentages for categorical data, and median and range for numerical data, were calculated to describe the study sample. The sampling scheme used in this study, in which affiliations and source articles are clusters, introduces multilevel dependency or correlation among the observations that can affect model parameter estimates. Therefore, we used a multilevel regression model for binary data to determine predictors of inaccurate citations. The model had a three-level data structure; the first level was citing articles, the second level was the source articles and the third level was affiliations. Statistical analyses were performed using the R environment for statistical computing (RRID:SCR_001905) (8) with the lme4 package (RRID:SCR_015654) (9). Significance level (alpha) was set at 0.05.

## Results

In total, we reviewed 4,535 citing articles (1,540 in the feasibility study and 2,995 in the verification set). The characteristics of these citing articles are shown in Table 3. The most common article types were original research (54.9%) and reviews (29.2%). The Vancouver or mixed citation style was used in most of the articles (92.0%). The median number of authors of the citing articles was five (range 1 to 65). The median impact factor of the 3,995 articles (88.1%) published in journals that had an impact factor at the time of publication was 3.262 (minimum 0.049, maximum 79.60). The source article was cited once (68.7%) in most cases. There were no discrepancies in the main characteristics between the feasibility study and verification set. The total number of citations of source articles was 2,526 and 4,912 in the feasibility study and verification set, respectively. The proportion of inaccurate citations in the feasibility study was 7.2% (183/2,526), while the proportion of articles containing at least one inaccurate citation was 11.1% (171/1,540). The presence of inaccurate citations was confirmed in the verification set, where the frequency of inaccurate citations was 10.3% (505/4,912), with the frequency of articles containing at least one inaccurate citation of 15.0% (449/2,995). Table 4 describes the types of citation inaccuracies in both sets. The most common finding was the citation of nonexistent findings (38.4%), followed by inaccurately cited numerical data/results (16.6%), inaccurate interpretation of findings (15.4%) and citations of quoted findings of another source (15.1%). The frequencies of the other types of inaccurate citations were below ten percent. In structured research articles, inaccurate citations mostly appeared in the introduction and discussion sections. Reviewers identified 13 chains of inaccurate citations in the feasibility study, in which the inaccurate citations appeared to have been copied from previous articles that had made the same citation error. These 13 chains included 44 articles with inaccurate citations (Figure 2), which accounted for approximately one fourth of all inaccurate citations (24%). The presence of chains of inaccurate citations was confirmed in the verification set, where 14 chains were identified, including 89 articles with inaccurate citations (Figure 2). Inaccurate citations included in the chains accounted for 19.3% of all inaccurate citations.

**Table 3.** Characteristics of citing articles

| Characteristics | Total<br>(n=4,535) | Feasibility study<br>(n=1,540) | Verification set<br>(n=2,995) |
| --- | --- | --- | --- |
| Article type,* n (%) |  |  |  |
| Original research | 2,490 (54.9) | 836 (54.3) | 1654 (55.2) |
| Review | 1324 (29.2) | 484 (31.4) | 840 (28.0) |
| Comment/Note | 54 (1.2) | 8 (0.5) | 46 (1.5) |
| Letter | 61 (1.3) | 15 (1.0) | 46 (1.5) |
| Brief/short report/communication | 50 (1.1) | 18 (1.2) | 32 (1.1) |
| Case report/series | 136 (3.0) | 59 (3.8) | 77 (2.6) |
| Study protocol | 19 (0.4) | 1 (0.1) | 18 (0.6) |
| Guidelines | 35 (0.8) | 10 (0.6) | 25 (0.8) |
| Pilot | 3 (0.1) | 2 (0.1) | 1 (0.1) |
| Opinion | 29 (0.6) | 0 (0) | 29 (1.0) |
| Editorial, editorial comment | 87 (1.9) | 11 (0.7) | 76 (2.5) |
| Other | 247 (5.4) | 96 (6.2) | 151 (5.0) |
| Self-citation, n (%) | 484 (10.7) | 92 (6.0) | 392 (13.1) |
| Citation style, n (%) |  |  |  |
| Vancouver or mixed | 4172 (92.0) | 1265 (82.1) | 2907 (97.1) |
| Harvard | 363 (8.0) | 275 (17.9) | 88 (2.9) |
| Number of authors, median (range) | 5 (1, 65) | 4 (1, 65) | 5 (1, 36) |
| Impact factor<sup>†</sup> |  |  |  |
| Have an impact factor, n (%) | 3,955 | 1,227 | 2,728 |
| Median (range) | 3.262<br>(0.049, 79.260) | 3.055<br>(0.051, 34.833) | 3.374<br>(0.049, 79.260) |
| Number of references in reference list |  |  |  |
| Median (range) | 41 (1, 1131) | 47 (1, 1131) | 38 (1, 620) |
| Time to citation, median (range), years | 6 (0, 24) | 6 (0, 23) | 5 (0, 24) |
| Number of citations, n (%) |  |  |  |
| 1 | 3,114 (68.7) | 1,046 (67.9) | 2,068 (69.0) |
| 2 | 775 (17.1) | 260 (16.9) | 515 (17.2) |
| 3 | 313 (6.9) | 121 (7.9) | 192 (6.4) |
| 4 | 149 (3.3) | 52 (3.4) | 97 (3.2) |
| ≥5 | 178 (3.9) | 60 (3.9) | 118 (3.9) |

\*Defined by the journal
<sup>†</sup>Retrieved from Journal Citation Reports for all journals that were indexed at the time when the citing article was published

**Table 4.**
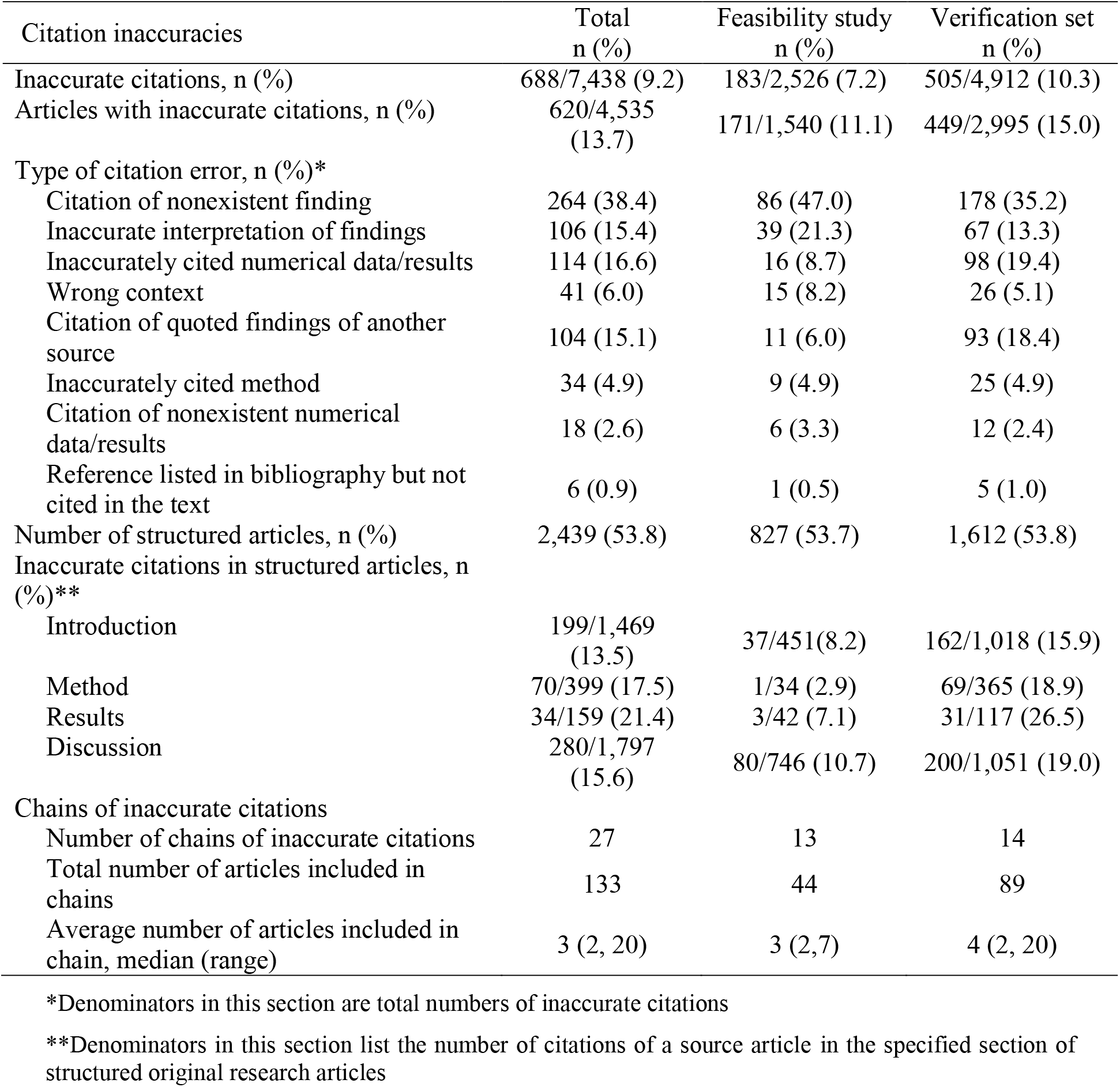
Citation inaccuracies

**Figure 2.**
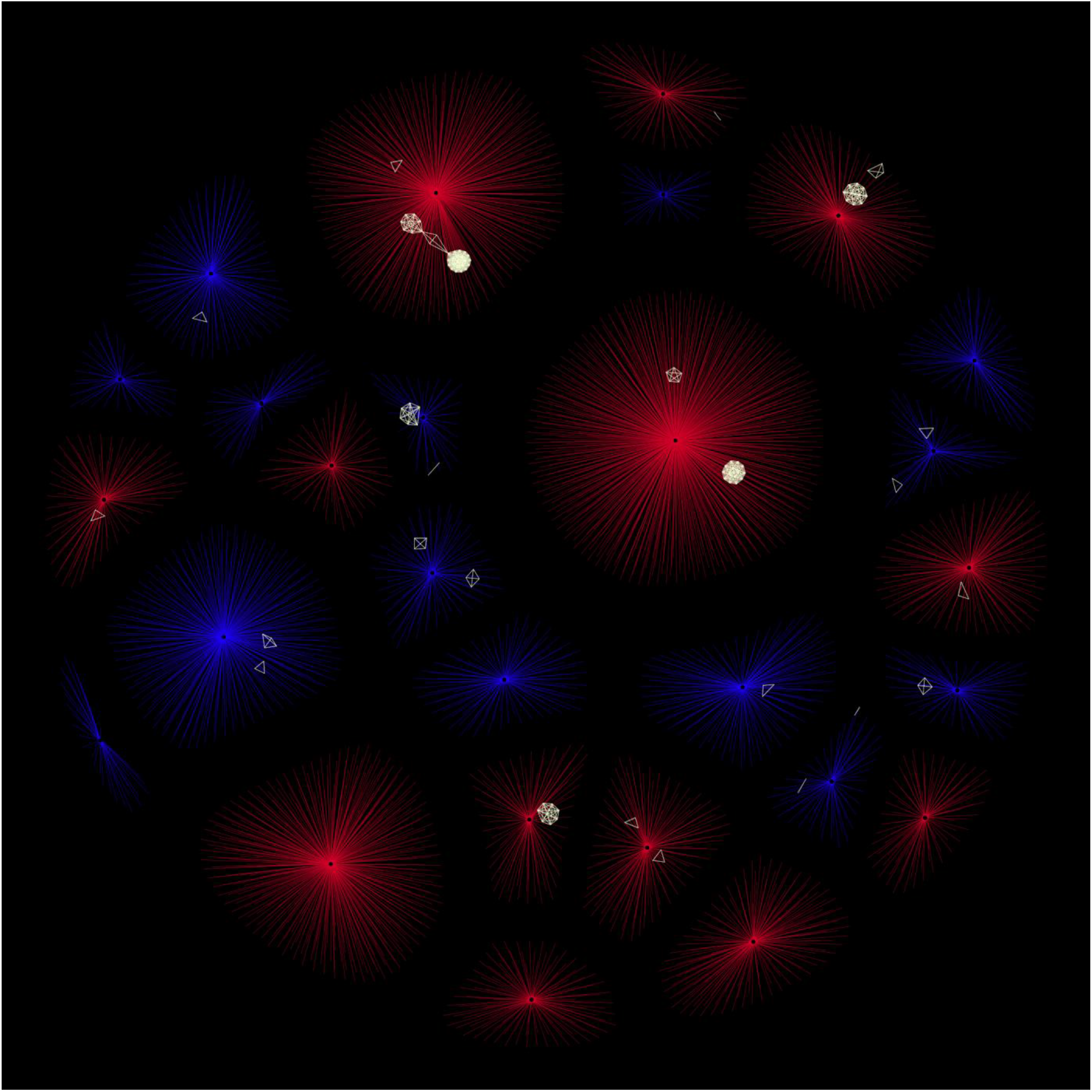
Presence of chains of inaccurate citations in the discovery and validation cohorts (circles - source articles, lines - citing articles, blue – feasibility study, red – verification set, white - chains of inaccurate citations)

Binary logistic regression models, with the presence of an inaccurate citation as a dependent variable in the model, are shown in Table 5. Statistically significant predictors in the univariate analyses were review articles, time elapsed time from publication to citation, impact factor and number of citations of the source article. Review articles, longer time elapsed from publication to citation, and a higher number of citations of the source article were associated with a greater risk of inaccurate citations in a multivariate model.

**Table 5.** Factors associated with inaccurate citations

| Independent variable | Univariate |  |  | Multivariate |  |  |
| --- | --- | --- | --- | --- | --- | --- |
|  | b | SE | p | b | SE | p |
| Review article | 0.22 | 0.09 | **0.023** | 0.22 | 0.09 | **0.022** |
| Time to citation (years) | 0.19 | 0.08 | **0.018** | 0.23 | 0.08 | **0.005** |
| Number of authors | -0.05 | 0.06 | 0.340 |  |  |  |
| Self-citation | 0.08 | 0.14 | 0.548 |  |  |  |
| Impact factor, Yes | -0.26 | 0.13 | **0.048** |  |  |  |
| Citation style, Vancouver or mixed | -0.14 | 0.15 | 0.373 |  |  |  |
| Number of citations of source article, >1 | 0.59 | 0.09 | **<0.001** | 0.60 | 0.09 | **<0.001** |
| Reference count | 0.11 | 0.06 | 0.057 |  |  |  |

Data were analyzed by multilevel regression models for binary data, with citation inaccuracy (yes vs. no) as the dependent variable.
b, regression coefficient; SE, standard error; p, p-value

Binary logistic regression models, with presence of chains of inaccurate citations as a dependent variable in the model, are shown in Table 6. Statistically significant predictors for the presence of chains in the univariate analyses were number of authors, self citation and number of references. In a multivariate model, higher number of references was associated with the occurrence of chains of inaccurate citations in biomedical literature.

**Table 6.** Factors associated with occurrence of chains of inaccurate citations

| Independent variable | Univariate |  |  | Multivariate |  |  |
| --- | --- | --- | --- | --- | --- | --- |
|  | b | SE | p | b | SE | p |
| Review article | 0.31 | 0.19 | 0.105 |  |  |  |
| Time to citation (years) | 0.30 | 0.17 | 0.073 |  |  |  |
| Number of authors | -0.09 | 0.00 | **<0.001** |  |  |  |
| Self-citation | -0.76 | 0.38 | **0.045** | -0.69 | 0.38 | 0.070 |
| Impact factor, Yes | -0.39 | 0.26 | 0.132 |  |  |  |
| Citation style | -0.20 | 0.29 | 0.483 |  |  |  |
| Number of citations of source article, >1 | -0.09 | 0.19 | 0.657 |  |  |  |
| Reference count | 0.39 | 0.12 | **<0.001** | 0.37 | 0.12 | **0.001** |

Data were analyzed by multilevel regression models for binary data, with citation inaccuracy (yes vs. no) as the dependent variable.
b, regression coefficient; SE, standard error; p, p-value

## Discussion

In this study, we found that inaccurate citations are common in biomedical scientific literature. At least one inaccurate citation was found in 11% of the reviewed articles in feasibility study. This finding was confirmed in the verification set of articles, where citation inaccuracies were detected in 15.0% of articles. The study was designed to determine the presence and types of inaccurate citations of the most cited original research articles from authors affiliated with two major research centers and to explore factors associated with inaccurate citations. The strengths of this study included collaboration with authors of the source articles to confirm and classify citation inaccuracies. Previous studies have used a “journal based approach” to determine the percentage of papers containing inaccurate citations (10). In contrast, we used a “source article based approach” to quantify the proportion of inaccurate citations for the most cited articles published by authors affiliated by our institutions. This approach yielded several important findings. Our results suggest that approximately one in ten citations of a highly cited article is inaccurate. Almost half of the citation inaccuracies in our sample were due to the citation of a non-existent finding, whereas 13.8% were due to an inaccurate interpretation of research findings. One fifth of the citation inaccuracies were due to chains of inaccurate citations, in which citation errors appeared to have been copied from previous papers. Review articles were more likely to contain inaccurate citations.

Although many studies have examined citation inaccuracies, our results may not be directly comparable due to differences in study design. Porrino et al. (11) used a similar approach to examine inaccuracies in citations of the Knirk and Jupiter (12) article, and found that 40% of citations were inaccurate by the time of the study (64/159). However, the generalizability of this finding was limited due to the fact that this study examined citations of a single article, which was selected because the authors were aware of the high rate of citation inaccuracies. Studies using the traditional journal based approach have reported that between 10% and 50% of papers contain citation inaccuracies (1,13). Only a few studies have reported a rate of inaccurate citations below 10% (14,15). One possible reason for this variability may be differences in the complexity (16) and scientific fields of the source articles, covering topics ranging from pure basic to applied clinical research.

In contrast to our results, previous studies have reported associations between citation inaccuracies and citation style (Harvard vs. Vancouver) (1), the number of authors (one vs. more than one) (17) or the number of references (18). These divergent results may also be partially due to study design differences. In contrast to other studies (14,19,20), we have found an association between inaccurate citations and journal impact factor. There were large differences in rates of inaccurate citations among our source articles (from 3.2% to 28.6%).

Our findings, along with previous studies demonstrating that citation inaccuracies are common, have several important implications for authors. Authors should adopt good citation practices, including those outlined in Table 7, when preparing manuscripts. These practices are important for all types of publications, including review articles, which were more likely to contain citation inaccuracies in our study. Inclusion of full texts of all citations in reference manager libraries should become prerequisite. Practices such as sharing libraries and asking multiple authors to check and confirm each citation may help to prevent common inaccuracies, including citations of non-existent findings and inaccurate interpretations of research findings.

**Table 7:**
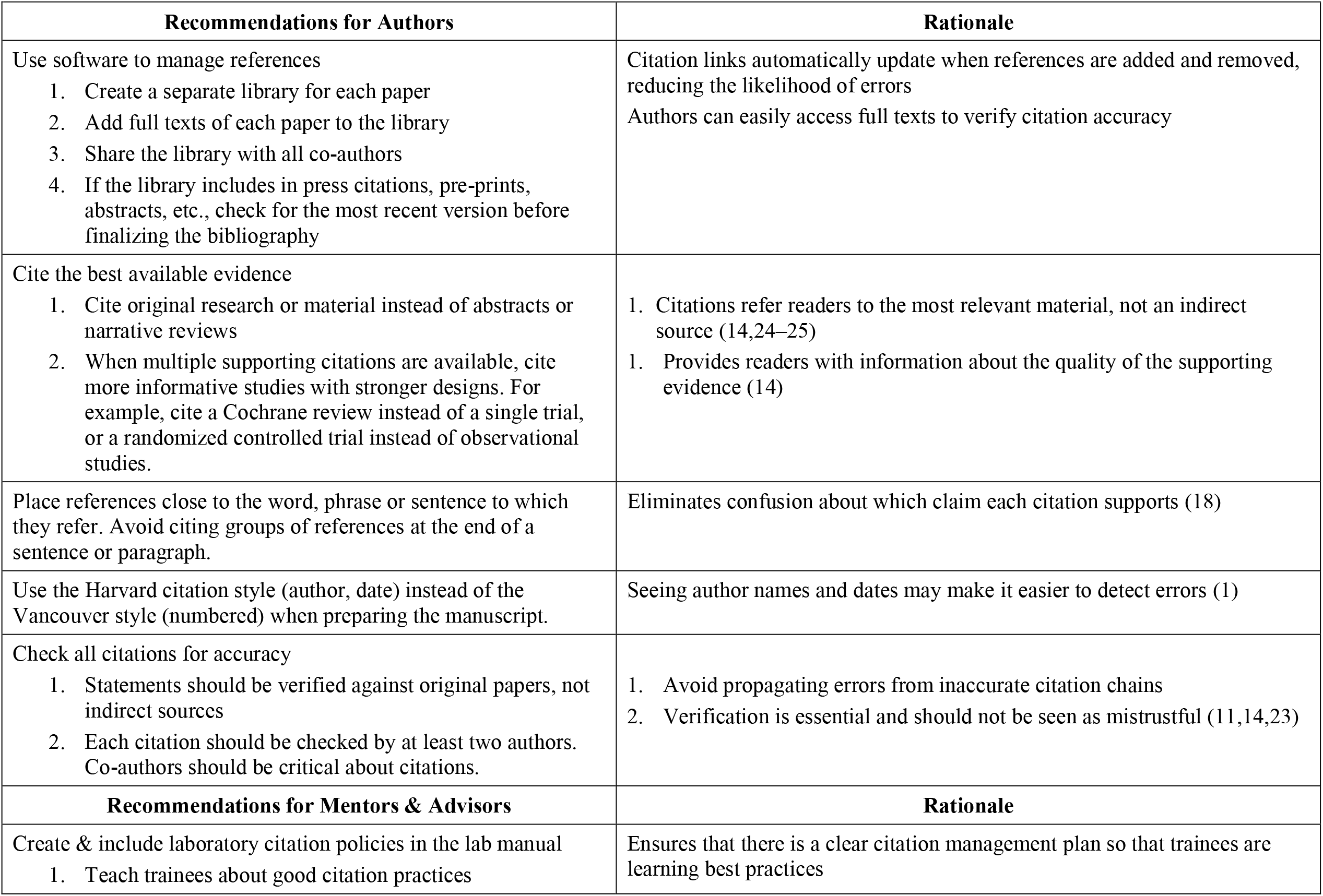
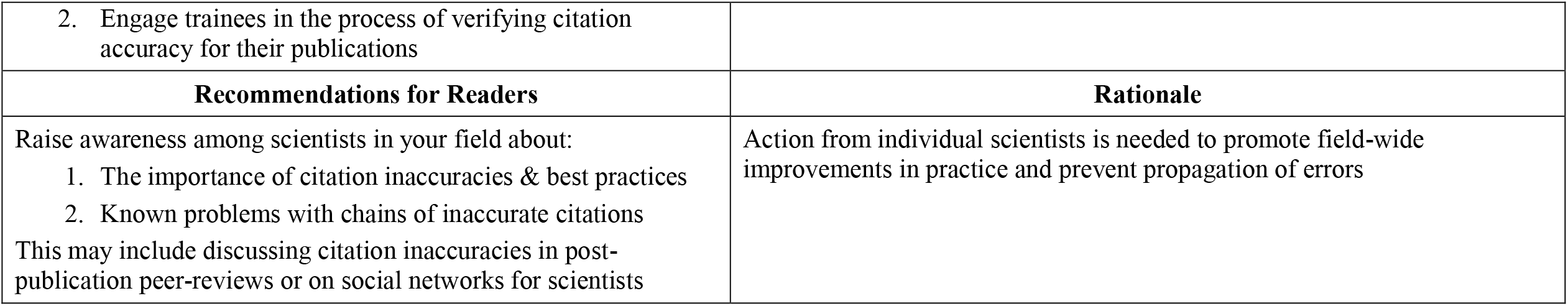
Actions authors, mentors and readers can take to encourage good citation practices and prevent errors

| **Recommendations for Authors** | **Rationale** |
| --- | --- |
| Use software to manage references<br>1. Create a separate library for each paper<br>2. Add full texts of each paper to the library<br>3. Share the library with all co-authors<br>4. If the library includes in press citations, pre-prints, abstracts, etc., check for the most recent version before finalizing the bibliography | Citation links automatically update when references are added and removed, reducing the likelihood of errors<br>Authors can easily access full texts to verify citation accuracy |
| Cite the best available evidence<br>1. Cite original research or material instead of abstracts or narrative reviews<br>2. When multiple supporting citations are available, cite more informative studies with stronger designs. For example, cite a Cochrane review instead of a single trial, or a randomized controlled trial instead of observational studies. | 1. Citations refer readers to the most relevant material, not an indirect source (14,24–25)<br>1. Provides readers with information about the quality of the supporting evidence (14) |
| Place references close to the word, phrase or sentence to which they refer. Avoid citing groups of references at the end of a sentence or paragraph. | Eliminates confusion about which claim each citation supports (18) |
| Use the Harvard citation style (author, date) instead of the Vancouver style (numbered) when preparing the manuscript. | Seeing author names and dates may make it easier to detect errors (1) |
| Check all citations for accuracy<br>1. Statements should be verified against original papers, not indirect sources<br>2. Each citation should be checked by at least two authors. Co-authors should be critical about citations. | 1. Avoid propagating errors from inaccurate citation chains<br>2. Verification is essential and should not be seen as mistrustful (11,14,23) |
| **Recommendations for Mentors & Advisors** | **Rationale** |
| Create & include laboratory citation policies in the lab manual<br>1. Teach trainees about good citation practices | Ensures that there is a clear citation management plan so that trainees are learning best practices |
| 2. Engage trainees in the process of verifying citation accuracy for their publications |  |
| **Recommendations for Readers** | **Rationale** |
| Raise awareness among scientists in your field about:<br>1. The importance of citation inaccuracies & best practices<br>2. Known problems with chains of inaccurate citations<br>This may include discussing citation inaccuracies in post-publication peer-reviews or on social networks for scientists | Action from individual scientists is needed to promote field-wide improvements in practice and prevent propagation of errors |

Scientists should also take steps to prevent the propagation of chains of inaccurate citations. These include carefully reading all papers prior to citation to prevent an inaccurate citation “domino effect,” not copying citations from other sources, and raising awareness about known chains of citation inaccuracies in the scientists’ fields. The practice of citing original papers without reading them has been already recognized in the literature as “lazy author syndrome” (21). While authors are responsible for ensuring that their citations are accurate, it is important to remember that most of the world’s scientific knowledge is still locked behind expensive paywalls. Universities spend millions every year on academic journal subscriptions for their students and faculty. While these costs may be manageable for some, they are prohibitive for many less wealthy scholars and institutions around the world. Scientists who are unable to access relevant articles may have to choose between not citing the reference, inappropriately citing an accessible secondary source, or citing the original article based on indirect information (i.e. the abstract or a citation in another paper). Teixeira et al. demonstrated that 15% of citations in ecology journals inappropriately referenced reviews instead of the original articles of authors who proposed the idea or reported research findings (22). Initiatives aimed at improving access to the scientific literature may help to address citation inaccuracies due to paywalls. These include pre-print servers, tools that locate open access versions of papers (i.e. UnPaywall), and funding agency policies that support or mandate open access publication.

Citation inaccuracies undermine the integrity of the scientific literature and can have serious consequences, however, good citation practices are rarely taught. Principal investigators can promote better practices by establishing standard citation protocols for their laboratories and engaging trainees in the process of verifying citation accuracy for their publications. The citations section of Table 7 includes references that provide more information regarding many of the practices described in the table. Other members of the scientific community can also develop incentives and implement strategies to improve citation accuracy. Table 8 provides an overview of strategies that journal editors can consider emphasizing the importance of citation accuracy and promoting good citation practices.

**Table 8:** Options journal editors can consider to encourage good citation practices

| Options for Journals | Rationale |
| --- | --- |
| The Instructions for Authors should include detailed guidelines, with recommendations for citing literature (for example, see “Recommendations for Authors” table) | Citation skills are rarely taught. Many authors are unaware of best practices. |
| Consider new policies and practices |  |
| 1. When submitting, ask authors to declare that they have checked all references for accuracy and have used primary references instead of indirect or secondary references | Emphasizes the need for good citation practices (26) |
| 2. Restrict the number of references | May make it easier for authors to maintain an overview of what they cite (14,27,28) |
| 3. Encourage editors and reviewers to check selected references | Random checks by editorial staff may remind authors about the importance of citation accuracy. Editors: (13,27); Reviewers: (17,29) |
| 4. Inform authors that citation accuracy is expected and checked | Shows that the editorial staff is committed to good citation practices (26) |
| 5. Institute a misquotations column to present cases of citation errors | Raise awareness of the consequences of inaccurate citations (28,30) |
| 6. Editors may consider whether certain statements really need one or more than one reference, particularly in discussion sections | Pressure to have a citation for every statement may increase the risk that authors include unnecessary or inappropriate citations (31) |

Limitation of our study is that source articles in the feasibility study and verification set of articles were each selected from one institution or department. This limitation only applies to the source articles because the citing articles came from different institutions and journals worldwide. However, findings were similar in both sets, suggesting that they may be generalizable to other institutions or departments. Selection of highly cited articles may limit the generalizability of the findings to articles with fewer citations. Additional limitations are the exclusion of articles not published in English and the use of a single–database based methodology.

## Acknowledgement

To the memory of Professor Goran Trajkovic, MD, PhD, Faculty of Medicine, University of Belgrade (1963-2019)

